# Single cell assessment of human stem cell derived mesolimbic models and their responses to substances of abuse

**DOI:** 10.1101/2023.05.12.540534

**Authors:** Thomas P. Rudibaugh, Ryan W. Tam, R. Chris Estridge, Albert J. Keung

## Abstract

The mesolimbic pathway connects ventral tegmental area dopaminergic neurons and striatal medium spiny neurons, playing a critical role in reward and stress behaviors. Exposure to substances of abuse during development and adulthood has been linked to adverse outcomes and molecular changes. The rise of human cell repositories and whole genome sequences enables human functional genomics ‘in a dish’, offering insights into human-specific responses to substances of abuse. Characterizations of *in vitro* models are necessary to ensure appropriate experimental designs and accurate interpretation of results. This study provides a comprehensive characterization of these models and their responses to substances of abuse, introducing new culture conditions for generating medium spiny neurons and dopaminergic neurons from human pluripotent stem cells. Single cell analysis reveals cell type-specific transcriptomic responses to dopamine, cocaine, and morphine, including compound and cell type-specific transcriptomic signatures related to neuroinflammation and alterations in signaling pathways. These findings offer a resource for future genomics studies leveraging human stem cell-derived models.

**Teaser:** Generation and characterization of a novel mesolimbic pathway model and its response to acute dopamine, morphine, and cocaine.

## Introduction

The mesolimbic “reward” pathway refers to the dopaminergic innervation of ventral tegmental area (VTA) dopaminergic neurons (DN) onto the medium spiny neurons (MSN) of the striatum. Signaling in this pathway regulates reward and stress behaviors, and pathway alterations either during development or adulthood are linked to a number of neurological disorders including schizophrenia and addiction^1–3^. Substances of abuse can act on this pathway through diverse mechanisms still being discovered including opioids or ethanol binding to µ-opioid receptors, and stimulants such as cocaine and amphetamines increasing the amount and duration of presynaptic dopamine impinging onto MSNs^4–10^. It is also possible and perhaps likely that these substances have additional impacts on neuronal and non-neuronal cell physiology, especially during early neurodevelopment. Exposure to these substances have been observed to drive many epigenetic, transcriptomic, structural, and synaptic changes both in adult MSNs and *in utero*^11–15^.

Our current understanding of cells of the mesolimbic pathway and their interactions with substances of abuse derives largely from rodent models and human epidemiological studies^16–19^. The rapid growth of human cell repositories, reprogramming technologies, and omics methods are driving considerable excitement for the potential of also performing human functional genomics in a dish. As an example, banked blood samples from hundreds to thousands of human donors can now be induced into pluripotent stem cells (hPSC) and then subsequently differentiated into target cell and organoid models. These models provide researchers the intriguing possibilities of performing experimental perturbations, identifying the effects of disease-linked mutations, and also capturing the magnitude and scale of effects driven by diverse human genetic backgrounds^20–35^. There is also increasing interest and needs for such models in preclinical testing^36^. However, these experimental models are still in their infancy with their utility limited by incomplete molecular and cellular characterization and variability in basal media compositions.

As a few example challenges related to mesolimbic cell types: multiple distinct models have been developed without clear comparisons of how they differ in cellular composition and responsiveness^23– 26,30–32,37–46^; most ventral forebrain organoids have not been characterized for their ability to generate MSNs^32,45^; it is unknown what the differences are between 2D and 3D models; it is a widely held, but not validated, assumption that 2D models are almost homogeneous systems especially compared to 3D organoids; it is unclear which cell types are driving responses measured in bulk to perturbations such as substances of abuse or microenvironmental factors; and finally, though not exclusively, as more complex systems are built from hPSC-derived cells, such as assembloids to model connections between brain regions, cell culture and media conditions will need to be engineered to be simultaneously compatible with the multiple neuronal or organoid systems being combined together. Therefore, it will be important to rigorously characterize, compare, and establish baseline datasets for these systems; these datasets will support the efficient and rigorous investment, design, and interpretation of expensive functional genomics experiments as well as of association studies.

Through single cell RNA sequencing (scRNAseq) and immunohistochemistry we quantitatively compare seven 2D and 3D models for their abilities to generate MSNs^23,26,30,32,44,45^. We also develop two novel models with the purpose of creating media conditions compatible with cocultures or assembloids. We then expose hPSC-derived organoids and assembloids to physiologically relevant doses of dopamine, cocaine, and morphine, perform scRNAseq, and identify cell-type specific transcriptional responses. This work provides new experimental frameworks and benchmarks for the use of hPSCs to model MSNs and DNs, provides direct comparisons between 2D and 3D systems, and reveals transcriptomic programs activated in these models in response to substances of abuse that may be relevant to long-term neuroplastic alterations observed in the mesolimbic pathway. These data and analyses serve as a resource for the design of future *in vitro* studies related to substance use disorders and *in utero* drug exposures, as well as to human stem cell derived models more generally.

## Results

### Development of striatal and midbrain organoids compatible with early assembloid generation

Our overall goals are to establish baseline comparisons of 2D and 3D models containing MSNs and DNs (with a focus on MSNs) by mapping their cellular compositions via scRNAseq, and to assess transcriptomic responses of a subset of these models to substances of abuses. However, we first take the opportunity to develop two new models that may fill existing gaps in contemporary experimental capabilities, with the goal that they can be included in our comparison of models.

Two key cell types of the mesolimbic pathway are MSNs expressing DARPP32 and DNs expressing tyrosine hydroxylase (TH). Both 2D neuronal and 3D cerebral organoid protocols have been regionally specified toward forebrain and midbrain lineages that are expected to include MSNs and DNs, respectively. However, while several 2D protocols have been shown to generate MSNs^23,26,30,43,44^ and DNs^31,39,40,46–51^, and 3D midbrain organoid protocols have been shown to generate DNs^24,25,27^, it is unclear if 3D forebrain organoids contain MSNs^32,45^. In addition, given the longer developmental timescales of organoid models, it would be useful in the future to be able to physically merge DN- and MSN-containing organoids into assembloids to simulate more complex neural pathways and projections. This is not currently possible as current forebrain and midbrain models are differentiated and cultured in different basal medias. With these two considerations in mind, our first goals are to develop one new organoid model, each, with forebrain and midbrain regional specifications that use media conditions mutually compatible with early fusion (at day 25). We also assess the expression of DARPP32 and other MSN markers in previously developed forebrain organoid models through immunostaining and RT-qPCR, prior to a comprehensive scRNAseq comparison of both the new and existing models.

Existing organoid models use a progression of basal media with varying compositions that are distinct from one another, for example in the balances of DMEM:F12 and neurobasal media, N2 supplement, and knockout serum replacement, among many other components. In addition, the lengths of patterning steps are highly variable between models and often extend well over four weeks. We also note that prior work generating forebrain organoids^32,45^ focused on the use of sonic hedgehog (SHH) to drive cells towards ventral fates deriving from the medial ganglionic eminence (MGE), yet striatal MSNs derive from the less ventral lateral ganglionic eminence (LGE) (Figure 1A-B). To address both of these issues, we begin with the whole brain organoid model developed by Lancaster and colleagues^34^ to provide a common basal media, as this media is the most ‘permissive’ of different regional specifications. We then layer on patterning factors informed by 2D differentiation models.

**Figure 1.**
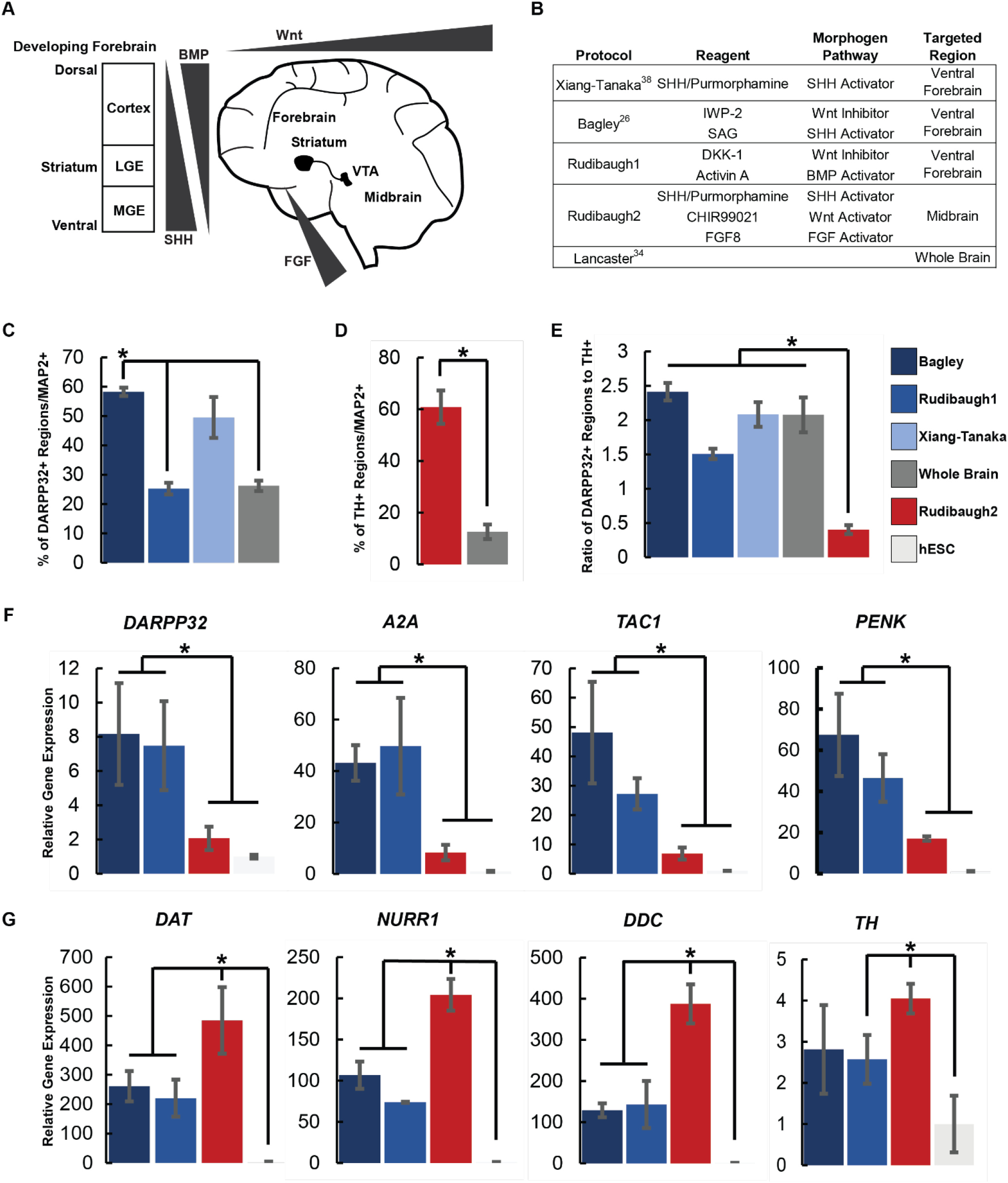
Development of striatal and midbrain organoids compatible with early assembloid generation. **(A)** Morphogen gradients associated with patterning the developing mesolimbic pathway. **(B)** Previously developed 3D forebrain organoid models and new forebrain and midbrain organoid models, with key patterning factors used, their indicated activities, and targeted regional specifications. **(C)** Immunostaining quantification of D90 forebrain and whole brain organoids. **(D)** Immunostaining quantification of D90 midbrain and whole brain organoids. **(E)** Immunostaining quantification of the ratio of DARPP32+ to TH+ regions in D90 forebrain, midbrain, and whole brain organoids. **(C-E)** Error bars represent 95% confidence intervals for n=4 biological replicates with 3 organoids per replicate. *p<0.05, one-way ANOVA with Tukey Kramer post hoc analysis. **(F-G)** RT-qPCR of D90 forebrain organoids, midbrain organoids, and hPSCs for **(F)** MSN markers *DARPP32, A2A, TAC1*, and *PENK* and **(G)** DN markers *TH, NURR1, DAT*, and *DDC*. Error bars represent 95% confidence intervals for n=5 replicates with 3 organoids per replicate. *p<0.05, one-way ANOVA with Tukey Kramer post hoc analysis.

Specifically, to target enrichment of forebrain striatal neurons, we begin with the basal media of the whole brain organoid model and add a combination of WNT inhibition by DKK-1 for anterior patterning, and moderate BMP activation by Activin A for medial dorso-ventral striatal development (Figure 1A-B, “Rudibaugh1”). This strategy draws upon prior derivations of 2D striatal neurons^30,52^. Five days after patterning is complete (day 30, D30), these organoids contain cells positive for the early striatal marker GSX2 and by D90 contain DARRP32+/CTIP2+ striatal neurons (Figures S1A-B). Among the three methods, the Bagley protocol produces the highest percentage of DARPP32+/MAP2+ neurons (Figure 1C). In addition, the Bagley organoids were found to be GSX2+ at D30 and DARPP32+/CTIP2+ at D90 (Figure S1C-D). The Rudibaugh1, Bagley, and Xiang-Tanaka models also express striatal markers *DARPP32, A2A, TAC1*, and *PENK*, at levels enriched over undifferentiated hPSCs as measured by RT-qPCR at D90 (Figures 1F, S1G-I).

To create midbrain organoids containing DNs, we again begin with the basal media of the whole brain organoid model and add a combination of SHH, WNT, and FGF pathway activators in line with previous 2D patterning methods^31^ (Figure 1A-B, “Rudibaugh2”). At D30, these organoids contain cells expressing the developing midbrain marker FOXA2^30,31^ and contain TH+/NURR1+ DNs (Figures 1D, S1E-I). These organoids also express DN markers *DAT, NURR1, DDC*, and *TH* at levels greater than Bagley and Rudibaugh1 organoids and undifferentiated hPSCs (D90, Figures 1G, S1H-I).

Reassuringly, the forebrain and midbrain organoids are all enriched for MSN and DN marker expression, respectively, relative to each other as well as to undifferentiated hPSCs (Figures 1F-G). Importantly, the patterning for both Rudibaugh1 and Rudibaugh2 models are complete by D25 at which point both organoid models could continue to mature in identical media and be fused to form an assembloid. We also note that the forebrain organoid model developed by Bagley and colleagues^32^ is also derived from the whole brain protocol^34^, and could be used in conjunction with the Rudibaugh2 model to create assembloids.

### scRNAseq reveals heterogeneous compositions and distinct enrichments of GABAergic subtypes in seven 3D and 2D striatal models

Immunostaining and RT-qPCR indicate two previously developed forebrain organoid models (Xiang-Tanaka and Bagley)^32,45^ do express striatal markers. Furthermore, the new forebrain (Rudibaugh1) and midbrain (Rudibaugh2) models, designed to share basal media components, contain striatal and dopaminergic cells, respectively. With confirmation that striatal neurons exist in the three 3D forebrain models (Xiang-Tanaka, Bagley, and Rudibaugh1)^32,45^, we next perform scRNAseq alongside four D45 hPSC-derived 2D striatal neuron models (Arber, Wu, Nicoleau, and Fjodorova)^23,26,30,44^, and ask how each model, and whether dimensionality, generally affects cell type composition (Figure 2A)^23,26,30,32,44,45^. We chose D90 3D organoids and D45 2D neural cultures as most publications cite these time points for stably matured and differentiated neuronal models.

**Figure 2.**
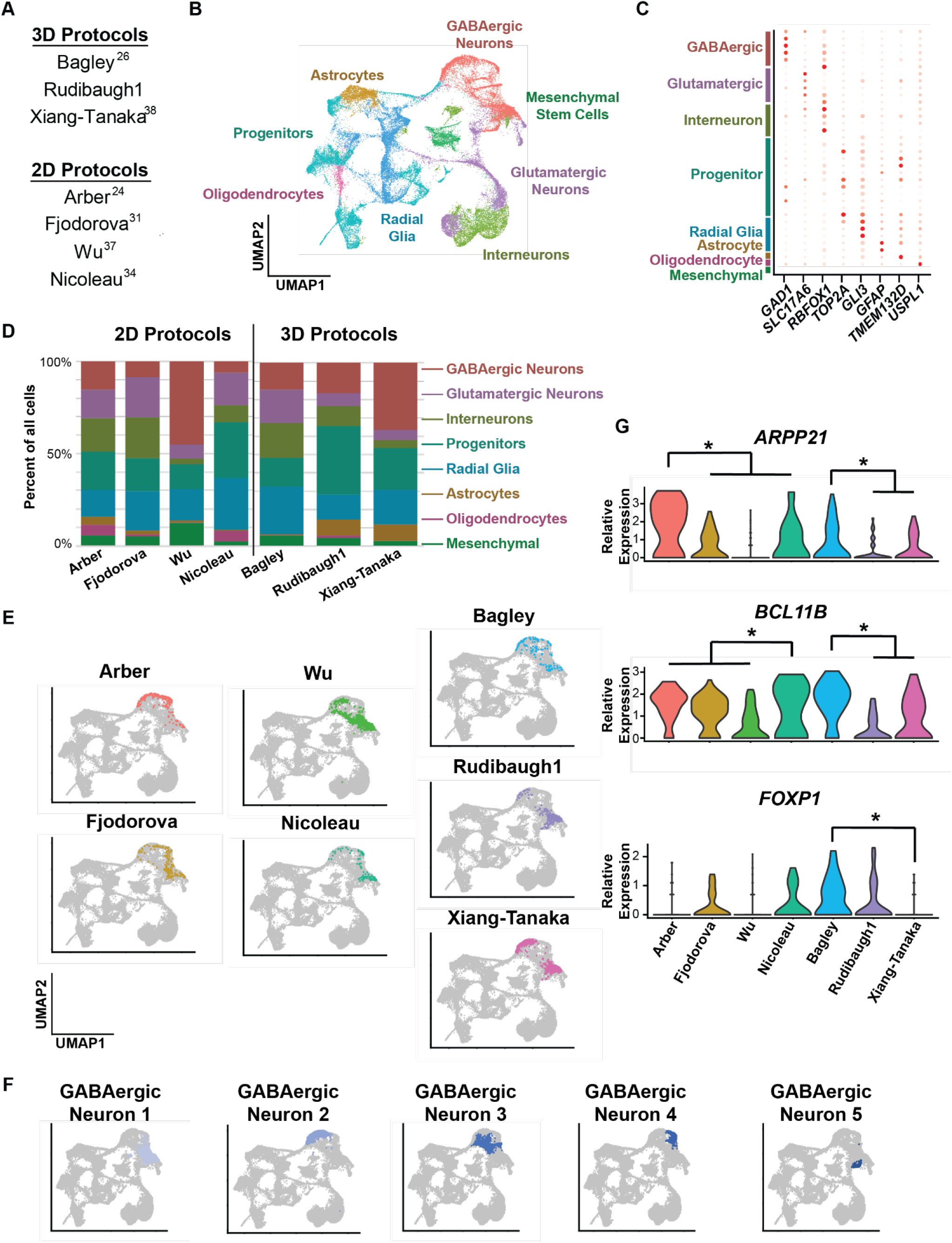
scRNAseq reveals heterogeneous compositions and distinct enrichments of GABAergic subtypes in seven 3D and 2D striatal models. **(A)** Table of the different striatal protocols used in the sequencing experiment. **(B)** UMAP plot shows positions of the 8 different identified cell types. **(C)** Dot plot of select marker genes showing how certain cell types were identified. **(D)** Cell type percentages of the 3 3D organoid protocols and 4 2D protocols show significant heterogeneity. **(E)** UMAP plots show the relative position of the GABAergic neurons in the seven different 2D and 3D culture models. **(F)** UMAP plots show the relative position of the five GABAergic neuron subclusters arranged from most to fewest number of cells. **(G)** Violin plots show GABAergic neurons in Bagley organoids have higher expression of striatal marker genes: *ARPP21, BCL11B (CTIP2)*, and *FOXP1* than the GABAergic neurons in the Xiang-Tanaka and Rudibaugh1 models. * false discovery rate q<0.05 relative to the other protocols.

Using combinations of marker genes as previously described^53^, we identify 34 distinct cell clusters comprising 8 broader cell classifications (Figures 2B-C, S2A). The broader cell classifications include *GAD1*+ GABAergic neurons, *SLC17A6+* glutamatergic neurons, *CNTNAP2+ or RBFOX1+* interneurons, *TOP2A+* progenitors, *GLI3+* radial glia, *GFAP+* astrocytes, *TMEM132D+* oligodendrocytes, and *USPL1+* mesenchymal stem cells. The specific genes associated with each classification are not the only ones defining each cell classification but are useful in presentation and discussion. As expected for organoid systems, the three striatal organoids show significant heterogeneity, with less than 40% of the cells being GABAergic neurons (Figure 2D). However, what is surprising is the status of the 2D models. 2D models are often presumed to be relatively homogeneous at this time point, or at least more so than what organoids achieve at any time point; yet they show a similar level of heterogeneity to the 3D models, with substantial percentages of GABAergic neurons, glutamatergic neurons, interneurons, oligodendrocytes, astrocytes, and stem and progenitor cells. In addition, there are no media components in the remaining maturation media that would be expected to direct remaining stem cells and progenitors to differentiate, or non-neuronal cells to be selected away, if the culture time were extended further.

We next narrow our analysis to ask if these models are enriched for specific subtypes of GABAergic neurons, seeking to identify a subcluster(s) that mimics MSNs of the striatum. There are five different subclusters of GABAergic neurons identified and each model generates a different proportion of each subcluster (Figure S2B). We observe enrichment of the GABAergic neuron 1 subcluster in the Rudibaugh1, Fjodorova, Wu, and Nicoleau models (Figures 2E-F, S2B). This subcluster expresses lower levels of GABAergic *GAD1* and *GAD2* and higher levels of glutamatergic *GRIN2A* and *GRM5* when compared to the other GABAergic neuron subclusters (Figure S2C). This suggests these are relatively immature neurons that express both GABAergic and Glutamatergic neuron markers, observed early in neurodevelopment as both neurotransmitters play a role in neurite outgrowth and axonal guidance^54,55^. The GABAergic neuron 2 subcluster is more abundant in the Xiang-Tanaka and Arber models; these cells exhibit relatively higher expression of neural progenitor markers *SOX4* and *SOX11*, suggesting these are relatively immature neurons as well^56^ (Figure S2B-C). GABAergic neuron subclusters 3 & 4, most abundant in the Bagley and Arber models, have robust expression of *GAD1* and *GAD2* along with higher expression of the striatal marker *ARPP21*, suggesting these clusters most closely recapitulate MSNs (Figure S2B-C). Interestingly, Rudbaugh1 is the only model generating GABAergic neuron subcluster 5 with as yet unclear physiological origins, characterized by higher levels of *ARPP21, LAMP5*, and *GRM5*. These results suggest the Bagley and Arber models have the highest percentages of MSN-like GABAergic neurons and may therefore be the best current 3D and 2D models respectively for modeling the striatum. This could be further tested by assessing the relative abundance of MSN marker genes in GABAergic neurons; however, due to limitations in scRNAseq sequencing depth it is challenging to detect the lowly expressed *DARPP32 (PPP1R1B), DRD1*, and *DRD2* transcripts^43^ (Figure S2D).

We therefore assess the expression of striatal neuron markers that are known to be more abundantly expressed: *ARPP21, CTIP2 (BCL11B)*, and *FOXP1* (Figure 2G). Reassuringly, GABAergic neurons in the Bagley model collectively express these genes at levels higher than the other 3D models (Figure 2G). The Arber model expresses elevated levels of *ARPP21*; however, it does not show higher expression of *BCL11B* and *FOXP1* relative to the other 2D models. Nicoleau and Fjodorova have relatively fewer cells in GABAergic neuron subclusters 3 and 4 but higher levels of *BCL11B* and *FOXP1*. Thus, it is difficult to conclude which 2D model generates the most MSN-like neurons mimicking the striatum. Overall, this analysis reveals that all current 3D and 2D models exhibit considerable heterogeneity, suggesting further work developing differentiation or sorting protocols is required if more homogeneous systems are desired; however, of those tested here, the Bagley model most closely mimics the striatum, and we therefore use it in the following studies with substances of abuse.

### Cell type specific differential gene expression responses to dopamine, cocaine, and morphine

Cerebral organoids could provide a compelling model system for studying transcriptional responses to substances of abuse due to the presence of multiple cell types present during development and their relative robustness in culture compared to 2D systems^22,35,57–60^. However, before they can be effectively used as *in vitro* models, further characterization is needed to establish baseline responses to these external stimuli^61,62^. We elect to characterize transcriptomic responses to dopamine, cocaine, and morphine here specifically because of their known relevance to the developing and mature striatum. In particular, MSNs are exposed to dopamine from a relatively young age^63^, and dopamine regulates cell cycle and proliferation in the developing ventral forebrain in E13 mice^63–65^ in addition to its role as a neurotransmitter. Cocaine’s effects in neurons are at least partly mediated through dopamine, driven by increasing the amount dopamine in synaptic clefts by blocking presynaptic reuptake through the dopamine transporter protein^66^. Morphine elicits responses in GABAergic neurons by binding to µ-opioid receptors^5^, while opioid receptors are also found on multiple other cell types in the developing brain including progenitors and glia^67,68^.

There are many experimental configurations possible to explore. We focus here on probing three: the effect of dopamine on forebrain organoids, as it is a primary presynaptic neurotransmitter for striatal MSNs; whether cocaine impacts MSNs in a manner distinct from direct dopamine treatment when in the presence of DNs in forebrain-midbrain assembloids, as the canonical understanding is that cocaine inhibits dopamine reuptake by DNs; and how morphine affects cells in forebrain organoids and assembloids. To address these questions, we expose Bagley forebrain organoids to physiologically relevant concentrations of dopamine or morphine, and Bagley-Rudibaugh2 forebrain-midbrain assembloids to cocaine and morphine (Figure 3A). We then perform an in-depth scRNAseq analysis and comparison of five cell types: GABAergic neurons, Glutamatergic neurons, Interneurons, Progenitors, and Radial Glia on their transcriptomic responses to all four conditions. Astrocytes, oligodendrocytes, and mesenchymal cells are excluded as no genes crossed the DEG threshold due to their low cell counts (Figures 3, S3, Supplemental Excel File 1).

**Figure 3.**
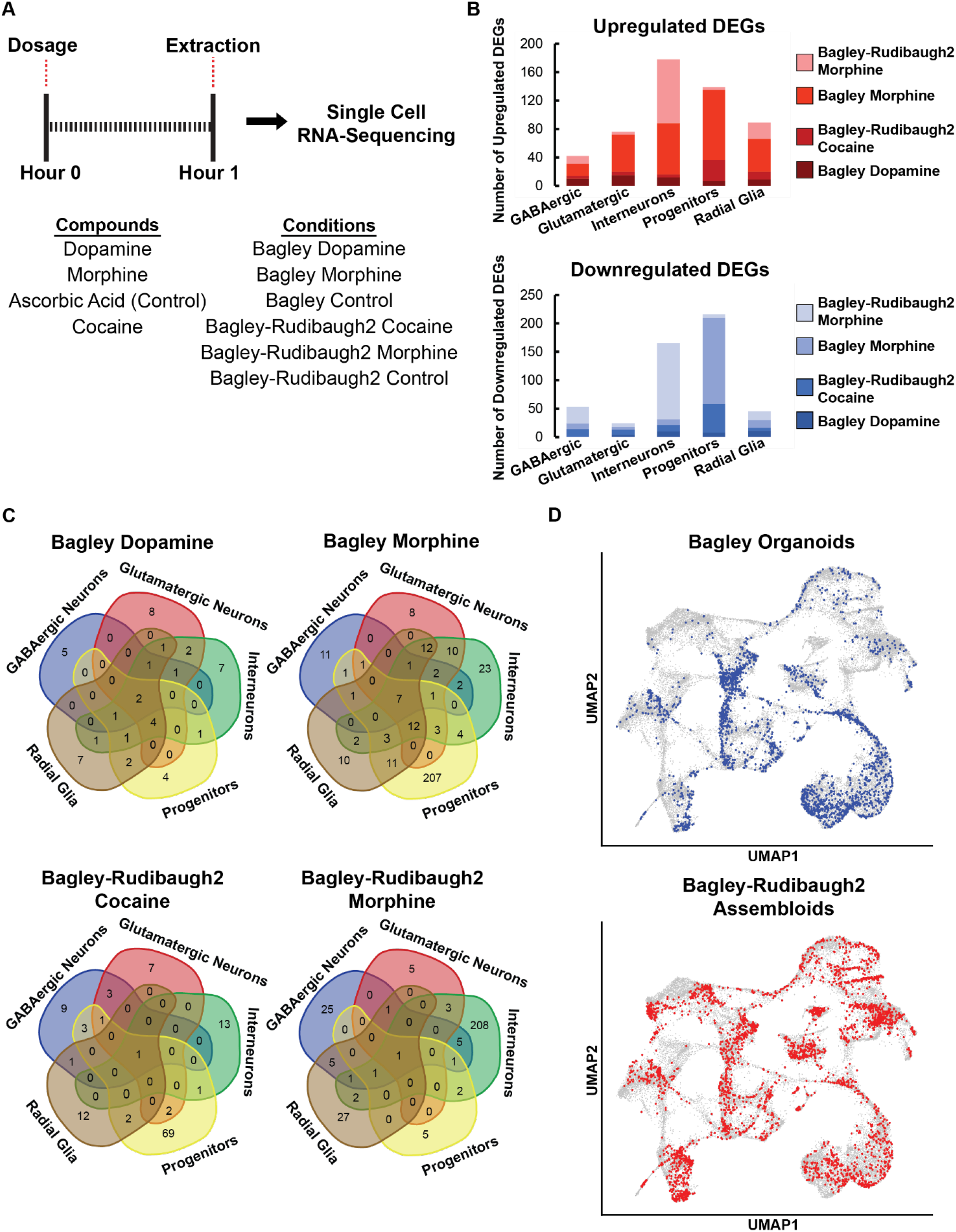
Cell type specific differential gene expression to dopamine, cocaine, and morphine. **(A)** Schematic of the six different conditions evaluated, the compounds used, and the duration of compound exposure. **(B)** Number of up and downregulated DEGs separated by cell type and condition. DEG defined as log2FC>|0.5|, expression in over 20% of sequenced cells, and false discovery rate q<0.05 relative to ascorbic acid dosed controls. **(C)** Venn diagrams comparing the DEGs across the 5 different cell types keeping the compound regimen constant. **(D)** UMAP plots show the relative positions of all cells from Bagley organoids and Bagley-Rudibaugh2 assembloids.

Our analysis of cell type specific differentially expressed genes (DEGs) relative to our ascorbic acid dosed vehicle controls reveal, somewhat surprisingly, that there are fewer DEGs in the GABAergic and Glutamatergic neurons compared to the Interneurons, Progenitors, and Radial Glia (Figures 3B, S3A). Thus, the effects of these compounds on transcriptomes may be more diverse than expected. Indeed, when visualizing the overlap of DEGs between cell types there are very few common DEGs, and this pattern holds for all compound treatments (Figure 3C, Supplemental Excel File 2). Similarly, when assessing each cell type and their responses to each compound, there are very few common DEGs between compound treatments (Figure S3B, Supplemental Excel File 2). These suggest that each compound has relatively distinct effects from each other and that different cell types respond distinctly to the same compounds as well. Interestingly, this distinctiveness is observed even between similar cells from Bagley organoids and Bagley-Rudibaugh2 assembloids in response to morphine (Figure S3C). This suggests the nature of the organoids/assembloids themselves influences the transcriptomic response. A broader composition comparison suggests different proportions of the eight unique cellular subtypes in Bagley organoids versus Bagley-Rudibaugh2 assembloids, which is not entirely unexpected (Figure 3D). In addition, cells from each of the five cell types, while generally clustered together on UMAP plots, exhibit some differences between the organoids and assembloids, and may explain their distinct responses to the tested compounds (Figure S3D). Ideally, we would perform subtype specific differential gene expression rather than identifying DEGs between the five broader cell type classes. However, we are unable to do this due to limitations in the number of cells in each individual cluster.

### Gene set enrichment analysis of cell type specific responses to dopamine, cocaine, and morphine

To next identify the nature of the cell type specific responses to each distinct compound, we perform gene set enrichment analysis (GSEA) on the cell type specific DEGs following drug exposure and identify the top pathways and upstream regulators (Figure S4, Supplemental Excel File 3). Several cell types show no or very few differentially expressed pathways or regulators following compound exposure, including many of the GABAergic neurons to all compounds and most of the cell types to dopamine. However, while each cell type seemingly shows a unique transcriptional response to each compound, there are some common broader trends. For example, among the 20-cell type specific analyses (5 cell types, 4 conditions each), several regulated pathways are identified multiple times including cellular responses to external stimuli along with oxidative phosphorylation and synapse formation (Figure 4A). The most common cellular response pathway is eIF2 signaling, followed by PI3K, CREB, and mTOR signaling. This trend applies to all five cell types, suggesting the immediate responses to the compounds work through similar pathways.

**Figure 4.**
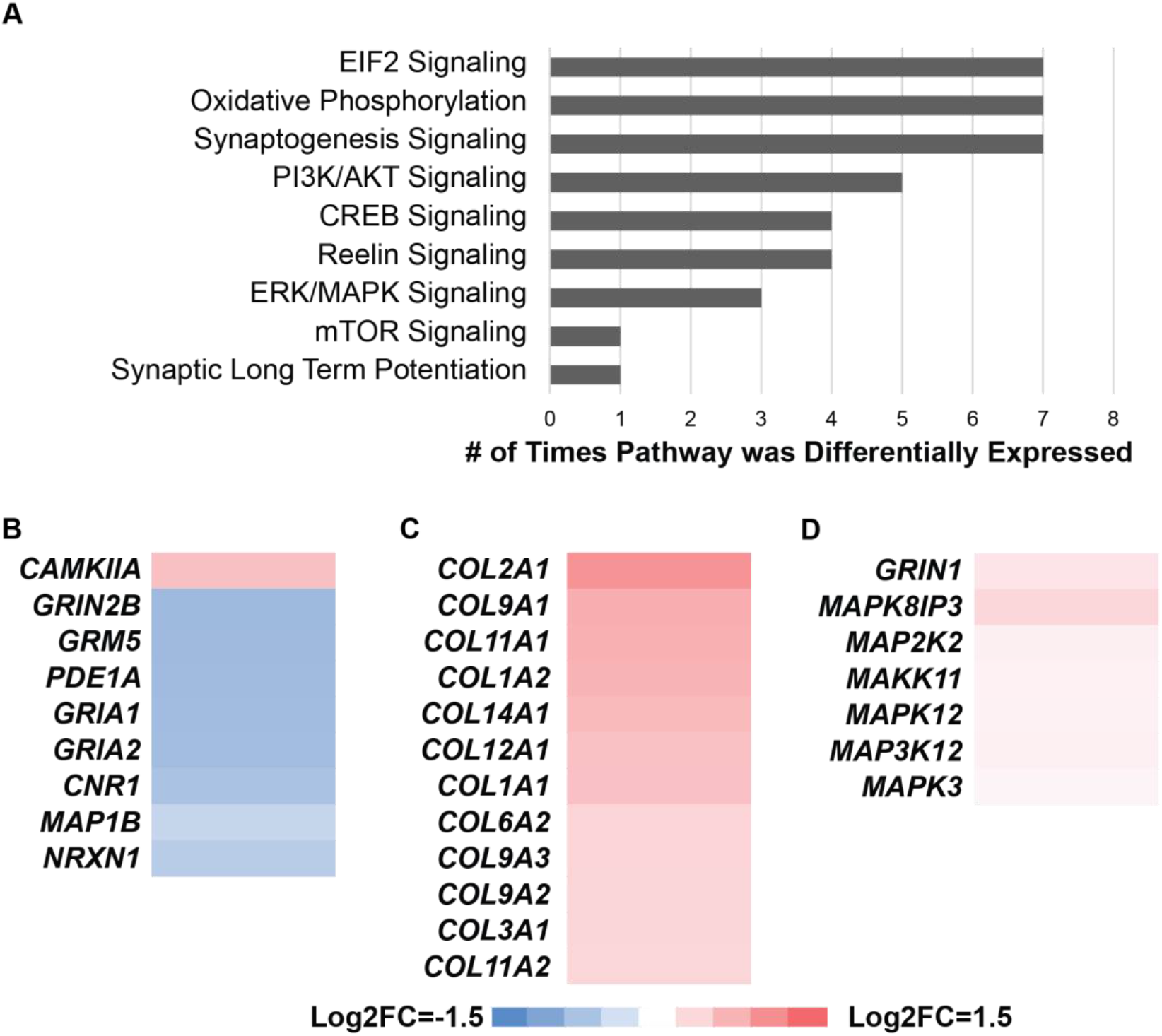
Gene set enrichment analysis of cell type specific responses to dopamine, cocaine, and morphine. **(A)** Counts of the top activated cell signaling and synapse related pathways based on the 20 unique cell-type specific DGE analyses performed. **(B-D)** Heat maps of differentially expressed genes from **(B)** Interneurons in Bagley-Rudibaugh2 assembloids exposed to morphine. **(C)** Progenitors in Bagley-Rudibaugh2 assembloids exposed to cocaine. **(D)** Glutamatergic neurons in Bagley organoids exposed to morphine. **(B-D)** All genes shown have false discovery rate q<0.05 relative to ascorbic acid dosed controls.

Thorough investigation of each of the 20-cell type-specific analyses reveals several responses that suggest a need for further investigation at a higher read depth. In this study, we highlight three transcriptional responses that exhibit a relatively higher number of differentially expressed genes (DEGs) and activate genes and pathways previously described in mice: the Interneuron response in Bagley-Rudibaugh2 assembloids to morphine, the Progenitor response in Bagley-Rudibaugh2 assembloids to cocaine exposure, and the response in glutamatergic neurons in Bagley organoids to morphine exposure. Interneurons in the Bagley-Rudibaugh2 assembloids exposed to morphine show downregulation of the synaptogenesis signaling and synaptic long term potentiation pathways (Figure S4). Within these pathways we observe downregulation of *CAMKIIA*, a gene previously linked with prenatal morphine exposure in mice; along with downregulation of multiple genes associated with *BDNF* expression, a gene similarly linked with prenatal morphine exposure^15,69,70^ (Figure 4B). Progenitors in Bagley-Rudibaugh2 assembloids exposed to cocaine exhibit activation of the stress response eIF2 and GP6 signaling pathways which regulate proliferation and cell cycle among others (Figure S4). In addition, multiple collagen genes are upregulated (Figure 4C). Collagen accumulation has not been previously linked with cocaine exposure, but the ECM plays an important, if understudied, role in neurodevelopment and in stress response and neuroplasticity^71^. Cocaine causes premature neural differentiation in early neural progenitors through oxidative stress mediated eIF2 signaling and it is possible this stress response also triggers increased expression of collagen proteins as a byproduct, although further testing is required to confirm this^72,73^. The glutamatergic neurons in Bagley organoids exposed to morphine exhibit upregulation of both the synaptogenesis and reelin signaling pathways; the top upregulated DEGs include *GRIN1* and multiple MAP kinases, suggesting a MAPK mediated pathway that may ultimately lead to regulation of synapse formation (Figure S4, Figure 4D)^74^. The increase in synaptic connections following opioid exposure is unique to glutamatergic neurons in mice models; suggesting organoids may be able to recapitulate this cell-type specific phenotype^69^. Overall, identification of cell type specific DEGs and pathway analysis reveals both some common trends and specific transcriptional responses to dopamine, cocaine, and morphine.

### Bulk analysis reveals roles for eIF2 and TGF-β signaling

A major limitation of scRNAseq is transcript capture efficiency and read depth per cell which limits the ability to identify cell type specific DEGs. However, the total reads from scRNAseq samples can also be aggregated and analyzed in bulk. This approach sacrifices single cell resolution in exchange for read depth and could provide a deeper understanding of how these models respond to each compound, responses that may be originally masked by the sparsity of scRNAseq data.

Taking this approach, the compound treated samples are compared to their vehicle controls in bulk (Figure 5, Figure S5). Among the top DEGs across all four treatment conditions: Bagley Dopamine, Bagley Morphine, Bagley-Rudibaugh2 Morphine, Bagley-Rudibaugh2 Cocaine, we observe multiple genes related to immune (*NFIA* and *NFIB*) and stress (*HSP90* and *NRFB1*) responses (Figure S5A-B), matching what is seen in the cell specific analysis. Acute drug exposure is known to alter neuroimmune responses *in vivo*, leading to cellular reactions including cytokine signaling, alterations in metabolic cycles, and even cellular apoptosis^75,76^. These phenotypes have been investigated both in mice and cell cultures; and recent work shows organoids can model neuroinflammation when exposed to acute methamphetamine; however, similar responses to cocaine and morphine have not been previously described in cerebral organoids^22,77^. Thus, we sought to identify what transcriptional pathways may be modulating this immune stress response. Using GSEA for all four treatment conditions, we observe upregulation of the aryl hydrocarbon receptor signaling pathway, which regulates the metabolism and transcriptional response to xenobiotics (Figure 5A).

**Figure 5.**
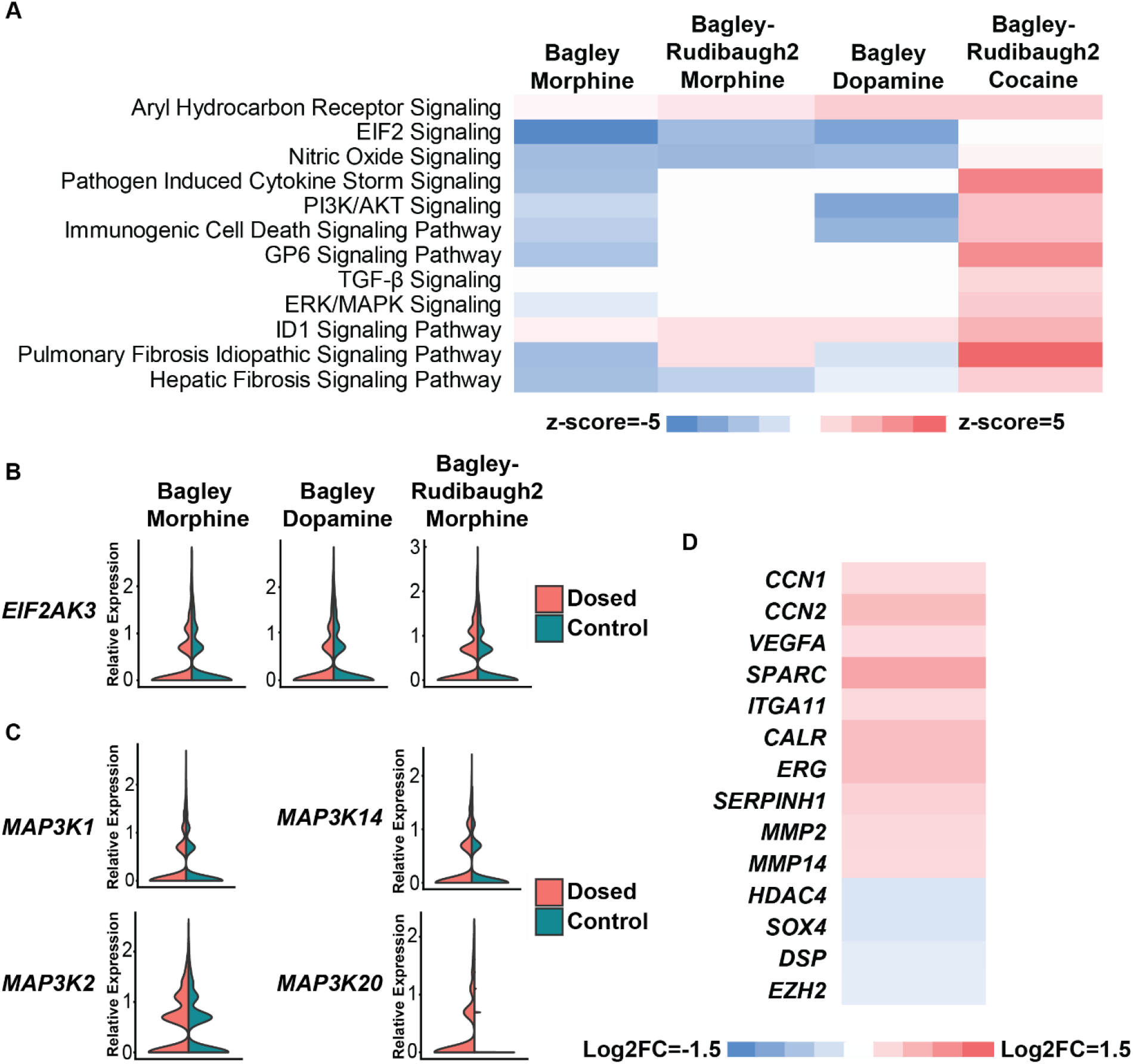
Bulk analysis reveals roles for eIF2 and TGF-β signaling pathways. **(A)** GSEA shows the Bagley Dopamine, Bagley Morphine, and Bagley-Rudibaugh2 morphine conditions exhibit downregulation of EIF2 Signaling. The Bagley-Rudibaugh2 Cocaine condition exhibits upregulation of TGF-β linked pathways. **(B)** The Bagley Dopamine, Bagley Morphine, and Bagley-Rudibaugh2 morphine conditions show upregulation of the *EIF2AK3* gene linked with ER stress and EIF2 phosphorylation. **(C)** Cocaine dosing causes upregulation of MAPKKK genes in Bagley-Rudibaugh2 assembloids. **(D)** Heat map of select, ECM-related DEGs following cocaine exposure. **(B-D)** All genes shown have log2FC>|0.5|, expression in over 20% of cells, and false discovery rate q<0.05 relative to ascorbic acid dosed controls.

Interestingly, the transcriptional response to cocaine was the most distinct. In the dopamine and morphine treated samples, we observe the downregulation of the eIF2 Signaling pathway, which is not downregulated in the cocaine treated organoids (Figure 5A). Instead, organoids exposed to cocaine exhibit multiple stress response pathways related to the cell cycle and proliferation including the ERK and AKT pathways, also previously linked to exposure to substances of abuse in the striatum^74,78,79^. Many of these pathways are linked to TGF-β signaling, which has been implicated in neurodevelopment and potentially responses to prenatal drug exposure^80–83^. In addition, we observe predicted upregulation of multiple fibrosis signaling pathways (Figure 5A). Central nervous system fibrosis has been reported following exposure to unknown pathogens; and results in an increase in ECM deposition^84^. This finding further supports a role for TGF-β signaling as it holds a critical role in regulating the ECM during neurodevelopment^81^. This also provides an explanation for the upregulation of collagen proteins observed in Progenitor cells exposed to cocaine (Figure 4C).

eIF2 functions as a stress signaling pathway which regulates global RNA translation, and previous work has demonstrated morphine and dopamine cause ROS mediated ER stress, ultimately leading to the expression of ER stress kinase gene *(PERK)*^85–87^. This in turn phosphorylates the eIF2α protein causing a cascading effect ultimately leading to a global reduction in RNA translation. We observe *PERK (EIF2AK3*) upregulated in the dopamine and morphine treated samples, suggesting they both contribute to ER stress leading to downregulation of eIF2 signaling (Figure 5B). In addition, all three conditions show the downregulation of multiple eIF2 pathway related genes including eiF2B, a known target for p-eif2α^88^; further supporting these compounds may be causing an ER-stress response acting through the eIF2 signaling pathway (Figure S5C). Taken together, this response suggests morphine exposure and potentially increased synaptic dopamine in the developing brain could lead to an immune regulation response and an ER regulated decrease in eIF2 Signaling. For cocaine exposure, further analysis reveals activation of multiple MAPKKK genes, indicating TGF-β could be potentially working through the ERK signaling pathway^74^ (Figure 5A,C). We also observe the upregulation of multiple ECM related genes including *SPARC, CCN1* and *CCN2* and the matrix metalloprotease genes *MMP2* and *MMP14* (Figure 5D), along with downregulation of genes linked with proliferation (*SOX4, DSP, HDAC4*, and *EZH2*). This suggests cocaine may affect the cytoarchitecture of developing neurons in cerebral organoids, and this response is potentially mediated by the TGF-β pathway^89,90^ (Figure 5D).

## Discussion

We have established hPSC differentiation protocols for generating striatal and VTA midbrain neurons by modifying the Lancaster whole brain organoid protocol^34^. The Wnt and BMP pathways were stimulated to make striatal organoids and the Wnt, SHH, and FGF pathways were stimulated to generate midbrain organoids. The resultant organoids express key markers of striatal and VTA neurons, respectively, at both D30 and D90. This demonstrates the whole brain organoid protocol can be modified to generate regionally specified organoids. These organoids share a common maturation media after D25, which allows for their fusion in the same media.

While, to our knowledge, this is the first time the presence of DARPP32+ striatal neurons have been confirmed in cerebral organoids, previous groups have generated ventral forebrain organoids, and there are numerous 2D protocols as well as a biomaterial-driven model^91^ for generating DARPP32+ striatal neurons. These models were developed by manipulating different morphogen concentrations to achieve neural differentiation, indicating there are multiple methods for generating striatal neurons *in vitro*. While each method generates some striatal neurons, and the literature often assumes a strong level of homogeneity in downstream analyses, our direct comparisons and composition analysis shows significant heterogeneity exists in all 2D and 3D protocols. Only a fraction of cells in all models are positively identified as GABAergic neurons. Only a fraction of those cells could be positively identified as striatal-like neurons, while other subtypes of GABAergic neurons express markers for relatively immature GABAergic neurons or markers for both GABAergic and Glutamatergic neurons. The remaining cells include several different cell types including undifferentiated progenitors, astrocytes, and radial glia. Together, this shows the composition differences between these model systems are relatively small, and none of these models exhibit a similar composition to the adult *in vivo* striatum but may better mimic the developing striatum. To increase the percentage of DARPP32+ neurons *in vitro* will require significant improvements in stem cell differentiation, or better methods for sorting out the neurons of interest without causing excessive cellular stress.

From another perspective, cellular heterogeneity within organoids can present advantages in efficiently assessing how different cell types respond to external stimuli and providing a more native microenvironment. Recent work has begun using organoid models to study responses to substances of abuse and demonstrated they are able to recapitulate several relevant phenotypes^22,57,60^. For example, Kindberg and colleagues recently demonstrated that cortical organoids exposed to cocaine exhibit an ROS driven cellular response that contributes to premature neural differentiation^92^. Our cell specific analysis also indicates each cell type exhibits differential gene expression following exposure to multiple substances of abuse. Reassuringly, GSEA of the cell type specific responses suggest many of the biological processes observed match what is seen in mouse models, including a neuroinflammatory response to morphine and downregulation of synaptogenesis and synaptic signaling in GABAergic neurons and interneurons following cocaine and morphine exposure (Figure S4, Figure 5A)^22,35,77,93^. Recent work has also suggested that cocaine causes an anti-inflammatory response in the striatum, and one role of TGF-β is an anti-inflammatory cytokine^94^. Cocaine may cause a similar response in cerebral organoids and represents another avenue for further investigation. Among the transcriptional responses not discussed in the results section are the Progenitor response to morphine in Bagley organoids and the GABAergic neuron response to morphine in both Bagley organoids and Bagley-Rudibaugh2 assembloids. DEGs in these cell types are linked with AMPA and NMDA receptors, both of which are linked with prenatal morphine exposure, suggesting another phenotype of interest, and possibly indicating these Progenitor cells are relatively immature neurons^15,76^.

We identified a novel role for TGF-β signaling in response to cocaine exposure, and for eIF2 signaling following morphine and dopamine exposure through bulk differential gene expression analysis. Both pathways hold critical roles in neurodevelopment, suggesting mechanisms by which prenatal drug exposure could have adverse effects on development. It could be interesting to further explore the apparent dissimilarity between the morphine treated assembloids from the morphine treated forebrain organoids. Both models induced down regulation of eIF2 signaling; however, the cell type specific DEGs show little overlap between the two organoid models, and indeed the morphine treated assembloids show higher DEG overlap with the cocaine treated assembloids (Figure S4B). One possible explanation is the slight compositional difference between Bagley organoids and Bagley-Rudibaugh2 assembloids. This also highlights the importance of conducting detailed pathway analysis when the overall read depth for cells is low, as individual gene expressions may exhibit variability. Altogether, this work provides novel insights into and baseline transcriptomic responses of hPSC derived cerebral organoids as brain region and cell type specific models for research on substance abuse and prenatal exposure.

## Materials and Methods

### hESC Cell Lines

H9 and H1 hESCs (WA09 and WA01; WiCell) were grown in E8 media (Thermo Fisher Scientific) in 6 well culture dishes (Greiner Bio-One) coated with 0.5µg/mL Vitronectin (Thermo Fisher Scientific). Cells were passaged every 3-5 days as necessary using 0.5mM EDTA (Thermo Fisher). All staining and qPCR experiments included in Figure 1 were carried out in H1 and H9 stem cells. Due to cost limitations, all sequencing experiments were limited to H1 stem cells.

### Organoid Cultures

The Xiang-Tanaka, Bagley, Monzel, an d Lancaster protocol organoids were grown as previously described^25,32,34,45^. The Rudibaugh midbrain and forebrain organoids were generated by modifying the Lancaster whole brain organoid protocol. Stem cells were grown to 75% confluency before dissociation into a single cell suspension using EDTA and Accutase (Fisher). 9,000 cells were plated into a low attachment U-bottomed 96 well plate(VWR) in hESC media supplemented with 50µM Y-27632(LC Labs) and 4ng/mL β-FGF(Thermo Fisher). hESC media contained DMEM-F12(Gibco), 20%v/v knockout serum replacement (Thermo Fisher), 3% v/v fetal bovine serum(Corning), 1% v/v MEM-NEAA(VWR), 1% v/v Glutamax(Gibco) and 7µL/L β-mercaptoethanol(Amresco). After 48 hours, half of the media was replaced with hESC media containing Y-27632 and β-FGF. After another 48 hours, half the media was again removed and replaced with hESC media without Y-27632 and β-FGF. On D6, organoid media was replaced with neural induction media supplemented with 50µg/mL heparin (Sigma). Bagley organoid media was further supplemented with 1µM of SAG(Sigma) and 2.5µM of IWP-2(Stemcell Technologies). Rudibaugh2 organoid media was supplemented with 100ng/mL SHH(VWR) and 2µM purmorphamine (VWR). Neural induction consisted of DMEM-F12, 1% v/v N2 supplement (Thermo Fisher), 1% v/v MEM-NEAA, and 1% v/v Glutamax. After 48 hours, 50% of the neural induction was replaced with fresh neural induction media. Bagley organoid media was again supplemented with heparin, IWP2, and SAG. Rudibaugh1 organoid media was supplemented with heparin; and Rudibaugh2 organoid media was supplemented with SHH and purmorphamine. On D10, neural induction was again changed with the same conditions as D8.

On D12, Bagley, Rudibaugh1, and Rudibaugh2 organoids were removed from the 96 well plates and embedded in Matrigel (Corning) as previously described^34^. Dimples were placed in Parafilm, and one organoid was transferred into each dimple using a cut 200µL pipet tip. 30µL of Matrigel was placed around the organoid and allowed to solidify at 37C for 30 minutes. The embedded organoids were then removed from the Parafilm and placed in a deep bottom 10cm plate (VWR) in COD – vitamin A media. The Rudibaugh forebrain organoid media was supplemented with 100ng/mL of DKK1(Peprotech), and the Rudibaugh midbrain organoid media was supplemented with 100ng/mL SHH, 2µM purmorphamine, and 3µM CHIR99021(Reagents Direct). COD – vitamin A media consisted of 50:50 DMEM-F12: Neurobasal (Gibco) mixture, 0.5% v/v N2 supplement, 1% v/v Glutamax, 0.5% v/v MEM-NEAA, 1% B27 supplement w/o vitamin A (Thermo Fisher), 1% v/v Penicillin/Streptomycin(VWR), 3.5µL/L β-mercaptoethanol, and 12.5µL/L 4mg/mL Insulin(). On D16, the media was replaced with COD + vitamin A media; which has the same components except the B27 w/o vitamin A is replaced with B27 supplement with vitamin A (Thermo Fisher) and placed on an orbital shaker (Thermo Fisher) rotating at 70rpm. The Rudibaugh forebrain organoid media was supplemented with 100ng/mL of DKK1 and the Rudibaugh midbrain organoid media was supplemented with 3µM CHIR99021 and 100ng/mL FGF-8(Peprotech). On D20, organoid media was replaced with fresh COD + vitamin A media. Rudibaugh1 organoid media was supplemented with 25ng/mL Activin A (R&D Systems) and Rudibaugh2 organoid media was supplemented with 100ng/mL FGF-8. On D24, media was changed, and Rudibaugh1 organoid media was supplemented with 25ng/mL Activin A. After D24, COD + vitamin A media was changed every 5-7 days as necessary until organoids reached desired timepoints.

### Assembloids

Organoid fusion assembloids were generated as previously described^32,45^. Briefly, one D25 midbrain and one D25 forebrain organoid were placed in a deep well V-bottomed 96 well plate (VWR) with 2mL of COD+vitamin A media; the plate was placed on the orbital shaker. The media was carefully changed after 48 hours, and after 96 hours organoid fusions were removed with a cut 1000µL pipette tip. Approximately 60% of organoids successfully formed fusion assembloids within 96 hours, those that were not successful were placed back in the V-bottomed plate and given another 96 hours to fuse. Assembloids were placed in 10cm plates and kept stationary for a further 48 hours before being placed back on the orbital shaker.

### Cryosectioning and Immunohistochemistry

Tissues were fixed in 4% paraformaldehyde (Sigma) for 15 minutes at 4C followed by 3, 10-minute PBS washes (Gibco). Tissues were placed in 30% sucrose overnight at 4C and then embedded in 10% gelatin/7.5% sucrose (Sigma). Embedded tissues were flash frozen in an isopentane (Sigma) bath between −50 and −30C and stored at -80C. Frozen blocks were cryosectioned (Thermo Fisher) to 30-µm. For immunohistochemistry, sections were blocked and permeabilized in 0.3% Triton X-100 (Sigma) and 5% normal donkey serum (VWR) in PBS. Sections were incubated with primary antibodies in 0.3% Triton X-100, 5% normal donkey serum in PBS overnight at 4C. Sections were then incubated with secondary antibodies in 0.3% Triton X-100, 5% normal donkey serum in PBS for 2 hours at RT, and nuclei were stained with 300nM DAPI (Invitrogen). Slides were mounted using ProLong Antifade Diamond (Thermo Fisher).

Images were taken using a Nikon AR confocal laser scanning microscope (Nikon). All samples within quantification experiments were imaged using the same laser intensity settings using the 10X objective. Quantifications were performed manually in FIJI. Organoid protocol names were first removed, and the images randomized before quantification to avoid bias. A 25×25 grid was overlaid over the image and each grid was manually counted as MAP2+ if over 75% of the region present had MAP2 axons. The images were then remeasured, and regions were considered DARPP32+ if over 50% of the MAP2+ cells appeared to be DARPP32+. For each condition, 4 independent replicates with 3 organoids per replicate were collected and measured.

### Real Time Quantitative PCR

Organoids were washed 3 times in ice cold PBS. Matrigel was dissolved by incubating the organoids in chilled Cell Recovery Solution (Corning) for 1 hour at 4C. The dissolved Matrigel was removed by rinsing 3 times in cold PBS. Total RNA was isolated using Direct-zol RNA MicroPrep Kit (Zymo Research) according to the manufacturer’s protocol. RNA samples were collected in 2mL RNAse-free tubes and chilled on ice throughout the procedure. cDNA synthesis was performed using 1µg of total RNA and the iScript Reverse Transcription Kit (BIO-RAD) according to the manufacturer’s protocol. Real time PCR was performed using the SYBR Green Supermix (BIO-RAD) according to the manufacturer’s protocol. Gene expression was compared to the reference genes GAPDH and ΔΔCq values were found by comparing against undifferentiated stem cell controls. For each qPCR condition, 5 independent biological replicates of 2-3 organoids per replicate were collected.

### Compound Treatment Experiments

D90 Bagley organoids and Bagley-Rudibaugh2 assembloids were treated with either a physiologically relevant dose of 1µM dopamine, 3µM cocaine (NIDA Drug supply program), or 13µM of morphine (NIDA Drug Supply Program), or an ascorbic acid control. Organoids were kept exposed to compounds for 1 hour before removal and immediate dissociation and fixation according to manufacturer’s protocols. 7-10 organoids were used for each condition.

### Dissociation and Library Preparation for Single Cell Sequencing

To obtain a single cell dissociation, the papain dissociation kit was used according to the manufacturer’s protocols (Worthington). Briefly, 2-3 organoids were initially broken into smaller pieces using a sterile razer before dissociation in papain solution. The cells were broken up further by gently pipetting using a 1000µL pipette.

Barcoding and library preparation of single cell suspensions was performed according to the manufacturer’s protocols (SplitBio). The cells were fixed and lysed, before multiple rounds of barcoding and cell measurements were employed to ensure the cells were in a uniform single cell suspension and each cell was given a unique barcode. Barcoded RNA sequences were shipped on dry ice before being sequenced on an Illumina HiSeq 4000×150bp at a depth of 50,000 reads per cell(Illumina Biosciences).

### Data Processing for Single Cell Sequencing

Raw sequencing data was preprocessed with splitp to pair oligo-dT primer and random hexamer barcodes used in the initial round of barcoding. A splici (spliced + intron) index was created using salmon (v1.7.0) using human GENCODE v39 and hg38 human reference genome. Sequences were mapped to the splici index using salmon and quantified using alevin fry (v0.5.0). Seurat (v4.1.1) was used to filter, process, and analyze gene expression matrices. The eight sequenced sublibraries were merged into a single Seurat object and reads not matching sample barcodes within a hamming distance of 1 were filtered out. QC filtering was performed removing cells that did not meet the following criteria: Count_RNA > 2000, nFeature_RNA > 1000, mitochondrial percentage < 10%, resulting in 43,844 high-quality cells for analysis. SCTransform was then used to normalize for read depth across cells, scale the data, and find variable features. The mitochondrial mapping percentage was regressed during normalization to remove this confounding source of variation. Principle component analysis (PCA) was performed and an ElboxPlot was generated to determine the number of informative dimensions (50). FindNeighbors, FindClusters, and UMAP dimensionality reduction visualization were performed. Clusters were identified based on the top 10 differentially expressed genes in each cluster, as well as analysis of known marker genes (replace this with however you did your cluster identification).

Cell types for each cluster were determined using previously identified marker genes^45,53,95^. All further downstream analyses were performed in RStudio using the high-performance learning computer available at North Carolina State University. Violin plots and dot plots were generated using open access R packages. The Ingenuity Pathway Analysis tool was used to perform the GSEA analyses presented. Differential gene expression was performed using the FindMarkers function on either all cells in the dosed or vehicle control or a specific cell subtype based on clustering identification. Genes were considered differentially expressed when they had a false discovery rate q<0.05, the gene was expressed in over 20% of the cells, and the Log2FC>|0.5|. Z-scores were obtained from the Ingenuity Pathway Software when only significantly differentially expressed genes were studied. Pathways and upstream regulators were considered up or downregulated when they had an adjusted p<0.05 and a z-score>|2.0|.

### Statistical Analysis

All error bars presented represent a 95% confidence interval assuming a normal distribution. P-values for immunostaining and qPCR experiments were calculated using a 1-way ANOVA test followed by a Tukey-Kramer test. False discovery rates for single cell sequencing were calculated using a Wilcox rank sum test. Information on the number of replicates and organoids per replicate can be found in each experimental procedures section.

## Acknowledgments

Cocaine and morphine were provided by the National Institutes of Drug Abuse (NIDA) as part of their drug supply program. We thank Dr. John Meitzen and Dr. Jeremy Simon for their help and thoughtful insights. We thank the In-Hyun Park research group for answering our questions and providing useful insights on their organoid protocol. We further thank the Park research group for kindly sharing their sequencing data with us and answering our questions. We thank Dr. Zuzana Drobna for providing the fluorescent cell line and all the assistance with our lab. We also thank Dr. Dilara Sen, Dr. Maria Fadri, and Sam Stuppy for their help and support.

## Author contributions

Conceptualization: TPR, RT, AJK

Wet Lab Experiments: TPR and RT

Upstream Bioinformatic Analysis: CE

Downstream Bioinformatic Analysis: CE, TPR, RT

Writing: TPR and AJK

## Data and materials availability

The sequencing data is available at: GSE225497.

