## Supplemental Information for "Single cell assessment of human stem cell derived mesolimbic models and their responses to substances of abuse"

Thomas P. Rudibaugh *et al.*

**This PDF file includes:**

Figures S1 to S5
Tables S1 to S3

**Other Supplementary Materials for this manuscript include the following:**

Excel Files S1 to S3

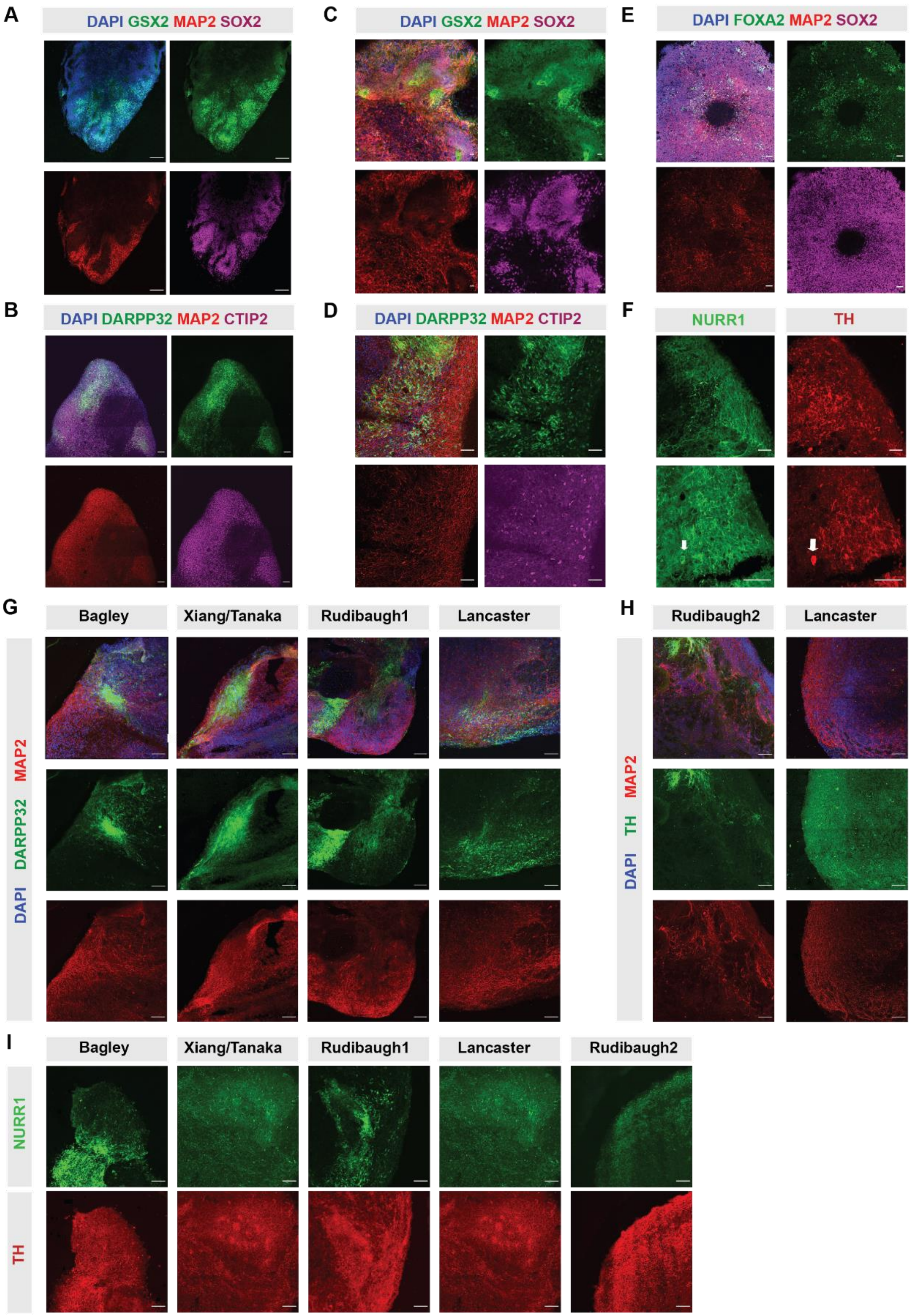
A
DAPI GSX2 MAP2 SOX2
B
DAPI DARPP32 MAP2 CTIP2
C
DAPI GSX2 MAP2 SOX2
D
DAPI DARPP32 MAP2 CTIP2
E
DAPI FOXA2 MAP2 SOX2
F
NURR1
TH
G
Bagley
Xiang/Tanaka
Rudibaugh1
Lancaster
DAPI DARPP32 MAP2
H
Rudibaugh2
Lancaster
DAPI TH MAP2
I
Bagley
Xiang/Tanaka
Rudibaugh1
Lancaster
Rudibaugh2
NURR1
TH

**Related to Figure 1; Figure S1:** ***Modification of the Lancaster whole brain model to generate forebrain and midbrain organoids***

**(A-B)** Immunostaining of **(A)** D30 and **(B)** D90 Rudibaugh1 forebrain organoids. **(C-D)** Immunostaining of **(C)** D30 and **(D)** D90 Bagley organoids. **(E)** Immunostaining of D30 Rudibaugh2 midbrain organoids. **(F)** D90 Rudibaugh2 organoids. **(G)** D90 Rudibaugh1, Bagley, and Xiang-Tanaka organoids. **(H)** D90 Rudibaugh2 organoids **(I)** D90 Bagley, Xiang-Tanaka, Rudibaugh1, Lancaster, and Rudibaugh2 organoids.

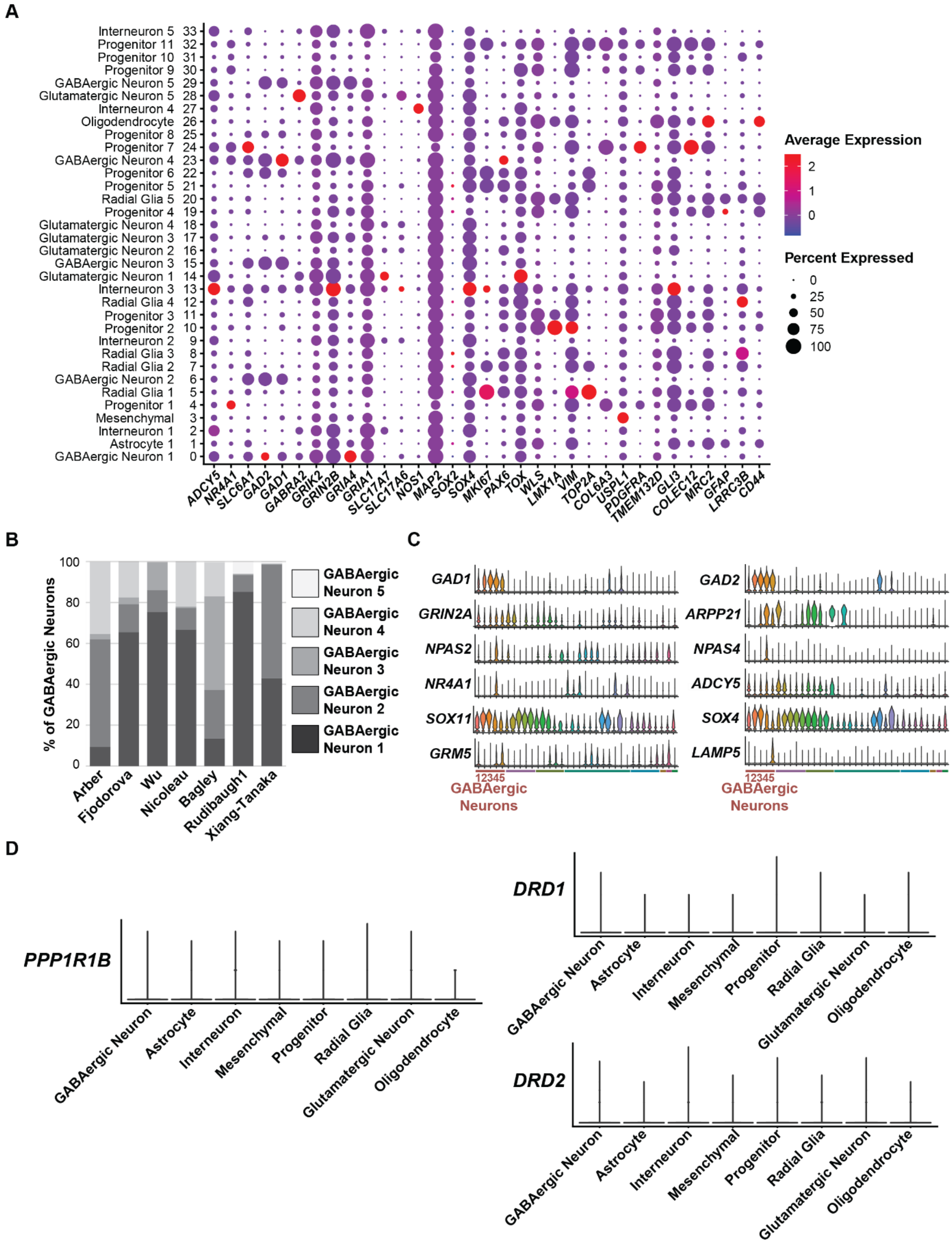
A
Interneuron 5 33
Progenitor 11 32
Progenitor 10 31
Progenitor 9 30
GABAergic Neuron 5 29
Glutamatergic Neuron 5 28
Interneuron 4 27
Oligodendrocyte 26
Progenitor 8 25
Progenitor 7 24
GABAergic Neuron 4 23
Progenitor 6 22
Progenitor 5 21
Radial Glia 5 20
Progenitor 4 19
Glutamatergic Neuron 4 18
Glutamatergic Neuron 3 17
Glutamatergic Neuron 2 16
GABAergic Neuron 3 15
Glutamatergic Neuron 1 14
Interneuron 3 13
Radial Glia 4 12
Progenitor 3 11
Progenitor 2 10
Interneuron 2 9
Radial Glia 3 8
Radial Glia 2 7
GABAergic Neuron 2 6
Radial Glia 1 5
Progenitor 1 4
Mesenchymal 3
Interneuron 1 2
Astrocyte 1 1
GABAergic Neuron 1 0
ADCY5 NR4A1 SLC6A1 GAD2 GAD1 GABRA2 GRIK2 GRIN2B GRIA4 GRIA1 SLC17A7 SLC17A6 NOS1 MAP2 SOX2 SOX4 MKI67 PAX6 TOX WLS LMX1A VIM TOP2A COL6A3 USPL1 PDGFRA TMEM132D GLI3 COLEC12 MRC2 GFAP LRRC3B CD44
Average Expression
2
1
0
Percent Expressed
0
25
50
75
100
B
% of GABAergic Neurons
100
80
60
40
20
0
Arber
Fjodorova
Wu
Nicoleau
Bagley
Rudibaugh1
Xiang-Tanaka
GABAergic Neuron 5
GABAergic Neuron 4
GABAergic Neuron 3
GABAergic Neuron 2
GABAergic Neuron 1
C
GAD1
GRIN2A
NPAS2
NR4A1
SOX11
GRM5
12345
GABAergic Neurons
GAD2
ARPP21
NPAS4
ADCY5
SOX4
LAMP5
12345
GABAergic Neurons
D
PPP1R1B
GABAergic Neuron
Astrocyte
Interneuron
Mesenchymal
Progenitor
Radial Glia
Glutamatergic Neuron
Oligodendrocyte
DRD1
GABAergic Neuron
Astrocyte
Interneuron
Mesenchymal
Progenitor
Radial Glia
Glutamatergic Neuron
Oligodendrocyte
DRD2
GABAergic Neuron
Astrocyte
Interneuron
Mesenchymal
Progenitor
Radial Glia
Glutamatergic Neuron
Oligodendrocyte

***Related to Figure 2; Figure S2: Clustering analysis reveals subpopulations of cells exhibit different abundances depending on the protocol***

**(A)** Dot plot of select marker genes for identifying the 8 different cell types. **(B)** Bar chart of the seven different models showing the relative abundance of the five different GABAergic neuron subtypes. **(C)** Clustered violin plots starting with the GABAergic neuron clusters on the far left showing expression of various genes of interest. **(D)** Violin plots depicting the expression levels of *DARPP32* (*PPP1R1B*), *DRD1*, and *DRD2* transcripts at 50,000 reads/cell show low expression in all cell types.

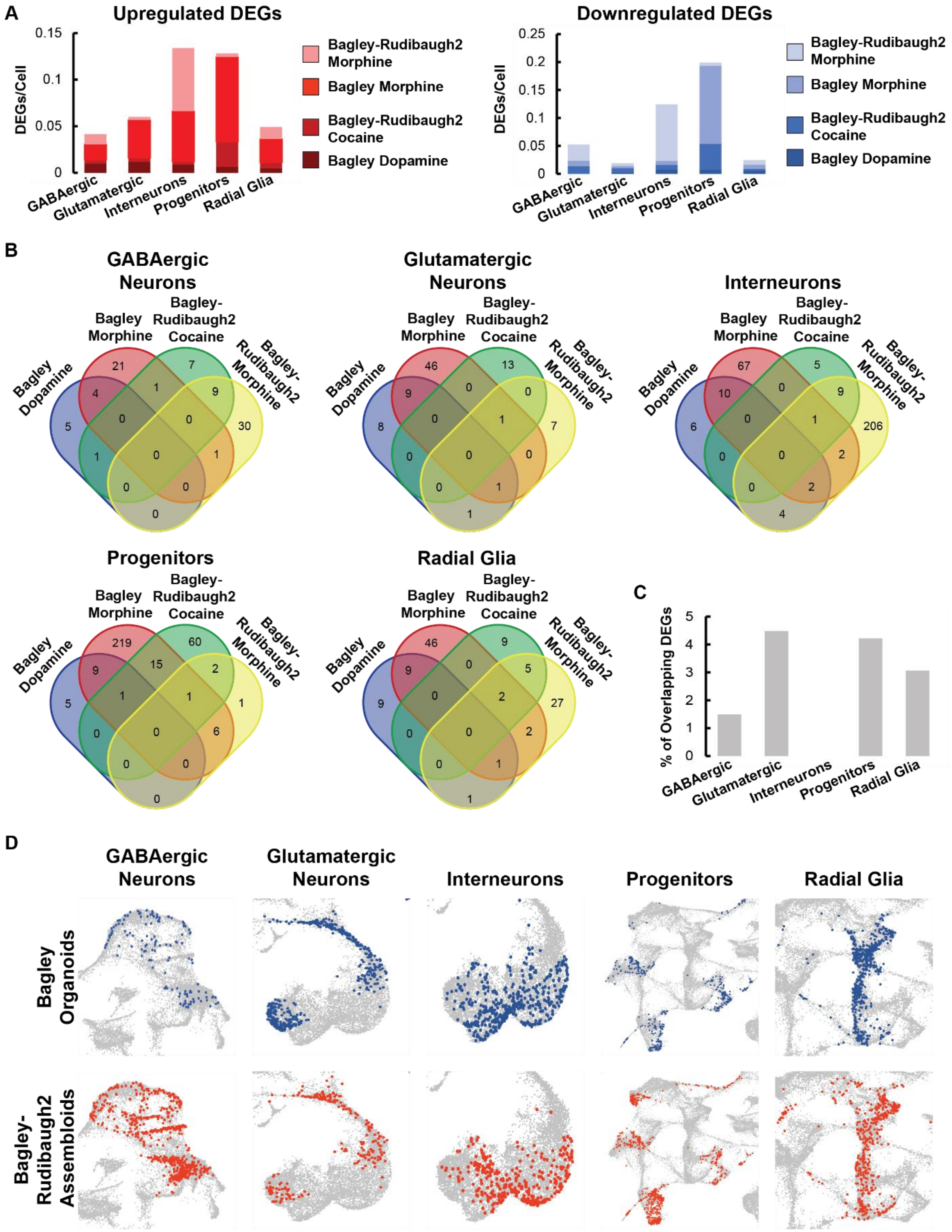
A
Upregulated DEGs
DEGs/Cell
Bagley-Rudibaugh2 Morphine
Bagley Morphine
Bagley-Rudibaugh2 Cocaine
Bagley Dopamine
GABAergic
Glutamatergic
Interneurons
Progenitors
Radial Glia
Downregulated DEGs
B
GABAergic Neurons
Glutamatergic Neurons
Interneurons
Progenitors
Radial Glia
Bagley Dopamine
Bagley Morphine
Bagley-Rudibaugh2 Cocaine
Bagley-Rudibaugh2 Morphine
C
% of Overlapping DEGs
D
Bagley Organoids
Bagley-Rudibaugh2 Assembloids

**Related to Figure 3; Figure S3:** ***Cell specific transcriptional analysis and comparisons following exposure to dopamine, cocaine, and morphine***
**(A)** The ratio of the total number of up and downregulated DEGs to the total number of cells in both control and dosed conditions. DEGs defined as log2FC >|0.5|, expression in over 20% of sequenced cells, and false discovery rate q<0.05 relative to ascorbic acid dosed controls. **(B)** Venn diagrams comparing the DEGs across the 4 different compound regimens keeping the cell type constant. **(C)** The ratio of overlapping DEGs between Bagley-morphine and Bagley-Rudibaugh2 morphine, when compared to the total number of DEGs while keeping the cell type constant. **(D)** UMAP plots showing the five different cell-types studied: GABAergic Neurons, Glutamatergic Neurons, Interneurons, Progenitors, and Radial Glia in Bagley Organoids and Bagley-Rudibaugh2 Assembloids

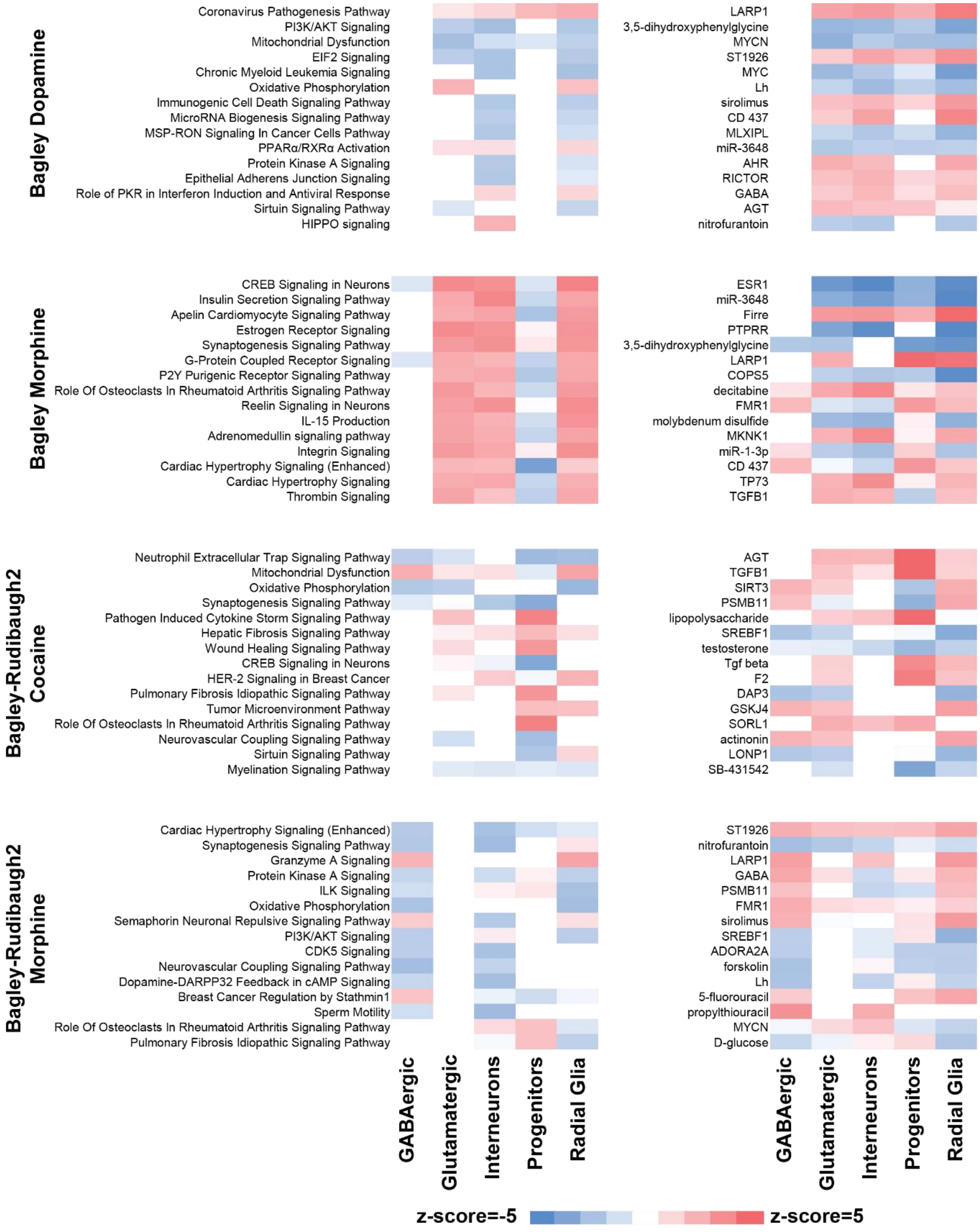

Bagley Dopamine
Coronavirus Pathogenesis Pathway
PI3K/AKT Signaling
Mitochondrial Dysfunction
EIF2 Signaling
Chronic Myeloid Leukemia Signaling
Oxidative Phosphorylation
Immunogenic Cell Death Signaling Pathway
MicroRNA Biogenesis Signaling Pathway
MSP-RON Signaling In Cancer Cells Pathway
PPARα/RXRα Activation
Protein Kinase A Signaling
Epithelial Adherens Junction Signaling
Role of PKR in Interferon Induction and Antiviral Response
Sirtuin Signaling Pathway
HIPPO signaling
LARP1
3,5-dihydroxyphenylglycine
MYCN
ST1926
MYC
Lh
sirolimus
CD 437
MLXIPL
miR-3648
AHR
RICTOR
GABA
AGT
nitrofurantoin
Bagley Morphine
CREB Signaling in Neurons
Insulin Secretion Signaling Pathway
Apelin Cardiomyocyte Signaling Pathway
Estrogen Receptor Signaling
Synaptogenesis Signaling Pathway
G-Protein Coupled Receptor Signaling
P2Y Purigenic Receptor Signaling Pathway
Role Of Osteoclasts In Rheumatoid Arthritis Signaling Pathway
Reelin Signaling in Neurons
IL-15 Production
Adrenomedullin signaling pathway
Integrin Signaling
Cardiac Hypertrophy Signaling (Enhanced)
Cardiac Hypertrophy Signaling
Thrombin Signaling
ESR1
miR-3648
Firre
PTPRR
3,5-dihydroxyphenylglycine
LARP1
COPS5
decitabine
FMR1
molybdenum disulfide
MKNK1
miR-1-3p
CD 437
TP73
TGFB1
Bagley-Rudibaugh2 Cocaine
Neutrophil Extracellular Trap Signaling Pathway
Mitochondrial Dysfunction
Oxidative Phosphorylation
Synaptogenesis Signaling Pathway
Pathogen Induced Cytokine Storm Signaling Pathway
Hepatic Fibrosis Signaling Pathway
Wound Healing Signaling Pathway
CREB Signaling in Neurons
HER-2 Signaling in Breast Cancer
Pulmonary Fibrosis Idiopathic Signaling Pathway
Tumor Microenvironment Pathway
Role Of Osteoclasts In Rheumatoid Arthritis Signaling Pathway
Neurovascular Coupling Signaling Pathway
Sirtuin Signaling Pathway
Myelination Signaling Pathway
AGT
TGFB1
SIRT3
PSMB11
lipopolysaccharide
SREBF1
testosterone
Tgf beta
F2
DAP3
GSKJ4
SORL1
actinonin
LONP1
SB-431542
Bagley-Rudibaugh2 Morphine
Cardiac Hypertrophy Signaling (Enhanced)
Synaptogenesis Signaling Pathway
Granzyme A Signaling
Protein Kinase A Signaling
ILK Signaling
Oxidative Phosphorylation
Semaphorin Neuronal Repulsive Signaling Pathway
PI3K/AKT Signaling
CDK5 Signaling
Neurovascular Coupling Signaling Pathway
Dopamine-DARPP32 Feedback in cAMP Signaling
Breast Cancer Regulation by Stathmin1
Sperm Motility
Role Of Osteoclasts In Rheumatoid Arthritis Signaling Pathway
Pulmonary Fibrosis Idiopathic Signaling Pathway
ST1926
nitrofurantoin
LARP1
GABA
PSMB11
FMR1
sirolimus
SREBF1
ADORA2A
forskolin
Lh
5-fluorouracil
propylthiouracil
MYCN
D-glucose
GABAergic
Glutamatergic
Interneurons
Progenitors
Radial Glia
GABAergic
Glutamatergic
Interneurons
Progenitors
Radial Glia
z-score=-5
z-score=5

***Related to Figure 4, Figure S4; Top regulated canonical pathways and upstream regulators based on GSEA***

GSEA of the cell type specific DEGs reveals the top 15 regulated canonical pathways and upstream regulators based on z-score. For a pathway or regulator to be significant z-score>|2|.

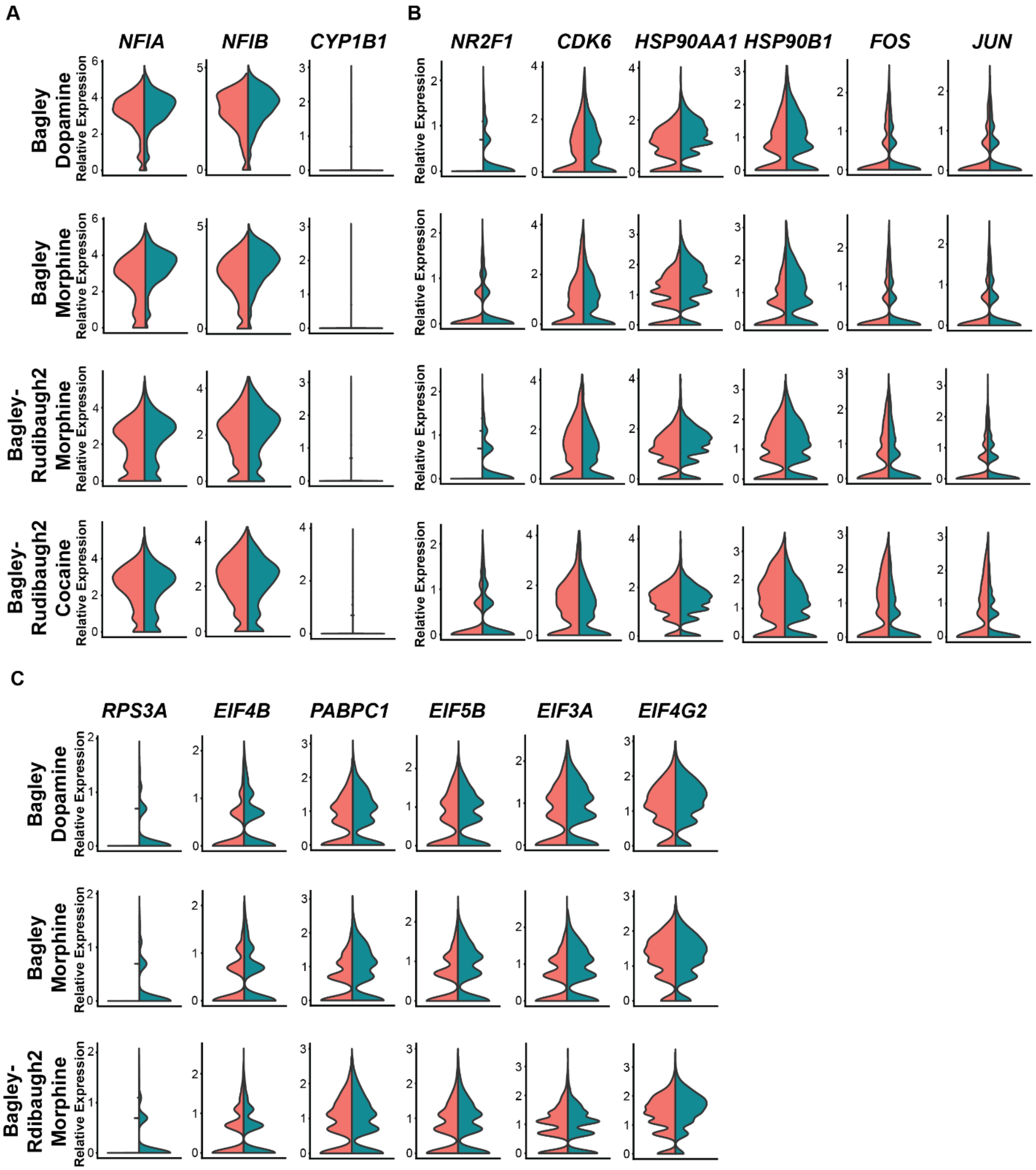
A
B
C
NFIA
NFIB
CYP1B1
NR2F1
CDK6
HSP90AA1
HSP90B1
FOS
JUN
RPS3A
EIF4B
PABPC1
EIF5B
EIF3A
EIF4G2
Bagley Dopamine
Bagley Morphine
Bagley-Rudibaugh2 Morphine
Bagley-Rudibaugh2 Cocaine
Bagley-Rdibaugh2 Morphine
Relative Expression

**Related to Figure 5; Figure S5:** ***Genes linked with neuroinflammation, eIF2α, and TGF-β are upregulated in compound dosed samples***

**(A)** Compound exposure causes differential expression of immune response genes in all four dosed conditions. **(B)** Compound exposure causes differential expression of stress response genes and heat shock proteins in all four compound regimens. **(C)** Downregulation of genes related to EIF2 signaling in Bagley Dopamine, Bagley Morphine, and Bagley-Rudibaugh2 morphine conditions. **(A-C)** All genes shown have false discovery rate $q<0.05$ relative to ascorbic acid dosed controls.

**Table S1**
**Related to Figure 1 and Figure S1, Primary Antibodies**

| Antigen | Species | Company | Catalog | RRID | Dilution |
|---|---|---|---|---|---|
| DARPP32 | Rabbit | Abcam | Ab40801 | AB_731843 | 1:50 |
| MAP2 | Mouse | Sigma-Aldrich | M1406 | AB_477171 | 1:250 |
| TH | Rabbit | Millipore | AB152 | AB_390204 | 1:300 |
| TH | Mouse | Immunostar | 22941 | AB_572268 | 1:250 |
| GSX2 | Rabbit | Millipore | Abn162 | AB_11203296 | 1:500 |
| SOX2 | Goat | R&D Systems | AF2018 | AB_355110 | 1:100 |
| CTIP2 | Rat | Abcam | Ab18465 | AB_2064130 | 1:250 |
| FOXA2 | Rabbit | Millipore | 07-633 | AB_390153 | 1:250 |
| NURR1 | Rabbit | Millipore | AB5778 | AB_92023 | 1:200 |
| GABA | Rabbit | Sigma-Aldrich | A2052 | AB_477652 | 1:200 |

**Table S2**
**Related to Figure 1 and Figure S1, Secondary Antibodies**

| Species | Flourophore | Company | Catalog | RRID | Dilution |
|---|---|---|---|---|---|
| Rabbit | 488 | Life Technologies | A21206 | AB_2535792 | 1:250 |
| Mouse | 546 | Thermofisher | A10036 | AB_2534012 | 1:250 |
| Rat | 647 | Jackson Immunoresearch | 712605150 | AB_2340693 | 1:125 |
| Goat | 647 | Jackson Immunoresearch | 705605003 | AB_2340436 | 1:125 |

**Table S3**
**Related to Figure 1 and Figure S1, qPCR Primers**

| Gene Symbol | Gene ID | Forward Primer | Reverse Primer |
|---|---|---|---|
| DARPP32 | 84152 | TTGGAAAATCCAGAAAACCG | CTGGTAGAAGCCGGTGAGAG |
| A2A | 135 | AGGCAGCAAGAACCTTTCAA | CTAAGGAGCTCCACGTCTGG |
| PENK | 5179 | GCTGTCCAAACCAGAGCTTC | TCTGGCTCCATGGGATAAAG |
| TAC1 | 6863 | TGGGGTTGAAAATTCAAAAAG | GGAGTTTCCTTCCTTTTCCG |
| TH | 7054 | TGTCTGAGGAGCCTGAGATTCG | GCTTGTCCTTGGCGTCACTG |
| DAT | 6531 | CAACAAGTTCACCAACTG | GGAGGAGAAGCTCGTCAG |
| NURR1 | 4929 | CAGGCGTTTTCGAGGAAAT | GAGACGCGGAGAACTCCTAA |
| DDC | 1644 | GGGGACCACAACATGCTGCTCC | AATGCACTGCCTGCGTAGGCTG |
| GAPDH | 2597 | ATGACATCAAGAAGGTGGTG | CATACCAGGAAATGAGCTTG |
